# Combining SMN2 splicing modifiers with HDAC6 inhibition greatly improves muscle function and survival in Spinal Muscular Atrophy

**DOI:** 10.1101/2025.01.15.633267

**Authors:** Alexis Osseni, Rasha Slika, Laurent Coudert, Agnès Conjard-Duplany, Laure Weill, Edwige Belotti, Zoé Clerc, Yann-Gaël Gangloff, Delphine Sapaly, Carole Vuillerot, Pascal Leblanc, Frédéric Charbonnier, Laurent Schaeffer

## Abstract

Spinal muscular atrophy (SMA) is a rare, progressive and severe neuromuscular disease. It is mostly caused by mutations in the SMN gene, which lead to the death of spinal cord motor neurons. In the absence of treatment, more than half of affected children die before the age of two. Recently, groundbreaking gene therapies were developed, allowing children to survive. However, a new clinical presentation of the disease has emerged in treated patients, characterized by ongoing functional deficits and a disability mainly due to persistent muscle atrophy. Over the last years, treatments of various animal models of neuromuscular disorders have shown the ability of inhibitors of the non-conventional histone deacetylase 6 (HDAC6) to reduce inflammation, fibrosis and muscle atrophy, and to ameliorate acetylcholine receptor distribution at the neuromuscular junction, microtubule network and mitochondrial transport in axons, indicating potential interest for the treatment of neuromuscular disorders. The present study was designed to properly characterize the effect of HDAC6 inhibition on muscle cells proliferation and differentiation and to evaluate *in vivo* if HDAC6 inhibition combined with the new standard of care treatments of SMA could ameliorate skeletal muscle and general status of a SMA mouse model. Here, we report that HDAC6 and tubulin acetylation controls myotube formation and maturation *in vitro*. In particular, HDAC6 inhibition increases the size of SMA patients-derived muscle primary myotubes. *In vivo*, when combined with ASOs inducing exon 7 inclusion in SMN2 RNA in motoneurons, HDAC6 systemic inhibition strongly improved muscle strength, mass, function, and longevity of the *Smn*^Δ7/Δ7^; hSMN2^+/−^ mouse model of SMA. These findings provide evidence that in muscle cells, HDAC6 is the only tubulin deacetylase and that selective inhibition of HDAC6 improves myogenic progression and that HDAC6 inhibitors are good candidates to ameliorate persisting symptoms of SMA patients treated with the new standard of care.

## Introduction

Spinal muscular atrophy (SMA) is a rare, genetic, progressive neuromuscular disease. SMA is classified into five subtypes (from 0 to 4), lower numbers corresponding to more severe forms. These subtypes are defined based on the highest motor milestone achieved and the age at which symptoms appear. Type 0 patients die perinatally, type 1 patients are characterized by the inability to sit and by a life expectancy usually not exceeding 2 years^1,2^. Half of the patients develop these particularly severe forms of the disease, while patients with the mildest forms remain ambulant throughout their lives despite a persistent muscle atrophy. In 95% of patients, SMA is caused by deletions or mutations in the survival of motor neuron 1 (*SMN1*) gene, leading to the degeneration of motor neurons in the spinal cord due to insufficient SMN protein levels^3^. The severity of the disease is mitigated by the expression of a *SMN1* paralogous gene named *SMN2*, which copy number in the genome increases as the severity decreases. *SMN2* genes mainly produce short-lived SMN protein due to the exclusion of exon 7 during splicing^4^. Accordingly, the therapeutic development in SMA mainly aimed at increasing SMN expression in patients. Besides the AAV-based gene therapy developed by Novartis (Zolgensma^@^), *SMN2* splicing-modifiers based on antisense oligonucleotides (ASO) or small molecules have been developed to increase exon 7 inclusion in the SMN2 RNA (respectively Nusinersen/Spinraza^@^ and Risdiplam/Evrisdy^@^)^5–8^. These disruptive treatments recently accessed the market and spare severe SMA patients from a certain death. However, fundamental and clinical data revealed that SMA is a developmental multiorgan disease, including severe intrinsic defects of skeletal muscles^9,10^. Importantly, muscle defects persist in treated SMA patients^11^ as well as in ASO-treated SMA-like mice of the Taiwanese model^12^. These data strongly suggest that additional therapies aimed at specifically improving muscle structure and function, which could be used in combination with the available SMN-targeting treatments, are critical for optimal care of SMA patients.

In this context, certain Histone deacetylases (HDACs) show promising characteristics. HDACs are post-transcriptional regulators that remove acetyl groups from lysine residues on histone and non-histone proteins. In humans, 18 HDACs are categorized into four classes based on their sequence homology to yeast HDACs: Class I (HDACs 1, 2, 3, and 8); Class II (HDACs 4, 5, 6, 7, 9, and 10); Class III/Sirtuins (SIRT 1–7); and Class IV (HDAC 11). Classes I, II, and IV are zinc-dependent enzymes with similar deacetylation mechanisms, while Class III HDACs rely on NAD+ to catalyze deacetylation. Most HDACs have several substrates and HDACs can have common substrates. In the case of microtubule deacetylation, α-tubulin lysine 40 deacetylation can be catalyzed by both the zinc-dependent HDAC6 and the NAD-dependent Sirt2^13,14^. HDAC6 is a non-conventional HDAC. It is strictly cytoplasmic and α-tubulin is its best characterized substrate^15^. During the past ten years, HDAC6 specific inhibitors have emerged as promising candidates for the treatment of cancer^16–20^ but also for neurological and muscular diseases such as, Amyotrophic Lateral Sclerosis, Charcot-Marie-Tooth’s, Alzheimer’s, Parkinson’s disease or Duchenne Muscular Dystrophy^21–26^. In a previous study, we have shown that HDAC6 interacts with the E3 ubiquitin ligase MAFbx/atrogin1 and that inhibiting HDAC6 expression protects against muscle atrophy in mouse^27^. Recently, we have evaluated the potential benefit of HDAC6 inhibition in a mouse model of DMD. This allowed us to identify HDAC6 as a new regulator of TGF-β signaling via the deacetylation of Smad3 regulating essential functions within cells by removing acetyl groups and serve as crucial mediators, responding to both normal physiological and disease-related signals^22^. In parallel, new Sirt2 inhibitors such as AGK2^28^ or SirReal2^29^ were developed. These selective Sirt2 inhibitors exhibit neuroprotective action in ischemic stroke by downregulating the expression of Akt/Foxo3a/MAPK pathways^30^. In the brain, these inhibitors attenuate lipopolysaccharides induced inflammation^31^.

In this report, we provide the first lines of evidence that HDAC6, but not Sirt2, controls the profile of tubulin acetylation in the skeletal muscle during myogenesis and that this acetylation activity impacts myotube formation and maturation. Moreover, we report here that HDAC6 inhibition in different SMA myoblast cell lines improved myogenesis rate, resulting in more numerous, more mature and better differentiated myotubes than in control cultures. Interestingly, these effects were independent of SMN expression in muscle. *In vivo*, pharmacological inhibition of HDAC6 combined with cerebrospinal injection of Nusinersen-like ASOs in a population of SMA-like mice from the Taiwanese model improved muscle mass and strength and increased longevity compared to mice injected with the ASO alone. Thus, taken together, these results suggest that HDAC6 inhibition could represent a valuable additional SMN-independent therapy to restore muscle function in SMA.

## Material and Methods

### Ethics statement, Animal models, treatments, and preparation of samples

All procedures using animals were approved by the Institutional ethics committee and followed the guidelines of the National Research Council Guide for the care and use of laboratory animals. Transgenic mild SMA-like males (SmnΔ7/Δ7; huSMN2+/+ 4 copies, FVB.Cg-Smn1tm1Hung Tg(SMN2)2Hung/J, strain #005058) were obtained from the Jackson Laboratory and were crossed with heterozygous Smn knock-out females (SmnΔ7/+), for generating 50% of severe SMA-like mice (FVB/NRj-SmnΔ7/Δ7; huSMN2−/+) and 50% of control mice (FVB/NRj-Smn+/Δ7; huSMN2−/+), both derived from the same strain after backcrossing on more than 4 generations with FVB/NRj female (Janvier Labs, Le Genest-Saint-Isle, France)^32^. All animals were housed in rooms at controlled temperature and humidity with ad libitum access to food and water. Mouse genotyping was performed by PCR assay using DNA extracted from tail biopsies. The experiments were performed blinded for genotype, treatment, and molecular analyses. All experimental mouse groups were randomly constituted with no pre-selection. All mice were injected at post-natal day 0 by single intracerebroventricular injection of 8μg of ASO 10-27 (ISIS-SMNRx; 5′-TCACTTTCATAATGCTGG-3′, Eurogentec) in 0.9% saline with 0.04% methylene blue, as previously described^33^. Following ASO, mice were divided in experimental groups receiving intraperitoneal injections of either Tubastatin A (TubA, APExBIO, #A4104; 10mg/kg/day) solubilized in DMSO and cyclodextrin ((2-hydroxypropyl)-βcyclodextrin, MW ca 1250-1480, H31133.14, Thermoscientific) according to manufacturer’s recommendations or saline solution supplemented with 4% DMSO and 38.4% cyclodextrin (vehicle), from P1 to P15 for histological and molecular analysis or from P1 to the age of reaching the ethical limit, for behavioral and survival curves. For all molecular and histological analysis, mice were euthanized at post-natal day 15 by pentobarbital administration (EXAGON ®) at 1 µL/g one hour after the last TubA or vehicle treatment. Then, muscles were dissected and either frozen in liquid nitrogen for protein and RNA extraction or embedded in optimal cutting temperature (OCT) medium (00411243, Q Path, VWR) and frozen in isopentane cooled with liquid nitrogen for cryostat sectioning. All the experiments were performed in the animal facility (BioMedTech Facilities, CNRS UMS 2009, INSERM US 36).

### Animal Behavioral analysis

The care and treatment of animals followed the national authority (French Ministry of Research and Technology) guidelines for the detention, use, and ethical treatment of laboratory animals. Grip strength test was performed in the forelimb of ASO-treated SMA-like mice co-treated with TubA or vehicles at P5, P9, P13, P17, P19 and P23. We measured the time during which the mice were able to sustain their own weight holding onto a thin metal rod suspended in air at 20 cm from a table. Five successive attempts were performed and separated by 30s resting period, with only the maximum value being recorded, according to the Treat-NMD guidelines (SOP (ID) Number SMA_M.2.1.002).

The righting reflex for ASO-treated SMA-like mice co-treated with TubA or vehicles was measured at P5 and P9. In this test, mice were placed in a supine position with all the four paws upright for maximum 40s. The time required for the mice to flip over onto the four paws back to a prone position was recorded. All mice were tested three times at each age. The behavioral test and analysis were based on previous reports^34^.

The ambulatory behavior was assessed in an open-field test for the two groups. The apparatus consists of a box measuring 15×15cm and divided into 25 squares of 3×3cm to test mice at 5 days of age, and a box measuring 28×28×5cm divided into 16 squares of 7×7 cm to test mice from 7 days. The squares adjacent to the walls were referred to as periphery, and the four remaining squares were referred to as center. The mice were tested individually and the field box was washed after each session. Each mouse was initially placed in the center of the arena and allowed to move freely for 5min. The number of peripheral and central square crossings were recorded manually.

### Cell culture

C2C12 cells was purchased at ATCC® (USA, #CRL-1772). Human immortalized culture of myoblasts from type 2 SMA patient was provided by Institute of Myology (Paris) under non-profit-to-non-profit uniform biological material transfer agreement (UBMTA). Human primary cells was provide by CBC Bio Tec of CRB unit of Hospices Civils of Lyon under non-profit-to-non-profit uniform biological material transfer agreement. Cells were seeded on matrigel-coated (matrigel® matrix, Corning) 35 mm diameter plates and were maintained as myoblasts in Dulbecco’s modified Eagle medium (DMEM) supplemented with 10% fetal bovine serum and 1% penicillin-streptomycin (Multicell). Then cells were differentiated in differentiation medium (DMEM medium supplemented with 2% horse serum, Bio Media Canada). Cells grown in 35 mm diameter plates were treated for either Western blot or immunofluorescence. For Western blot, cells were collected by trypsinization, washed with PBS, centrifuged, and stored at −20°C until used. For immunofluorescence, cells were fixed for 20 min in PBS-4% paraformaldehyde at room temperature, washed in PBS and stored at 4°C until used.

### Drug, antibodies, plasmids and shRNA and other reagents

Cells were treated with different drugs: tubastatin A (TubA at 5 µM, APExBIO, #A4101), tubacin (TBC, from 5 to 10 µM, Sigma, #SML0065), N-hydroxy-4-(2-[(2-hydroxyethyl)(phenyl)amino]-2-oxoethyl)benzamide (HPOB, at 5 µM APExBIO, #B4890), trichostatin A (TSA, at 10 µM, Calbiochem, #647), SirtReal2 (from 50 to 100 µM, Sigma, #) and AGK2 (from 0,1 to 50 µM, Sigma, #SML1514). All drugs stock solution were dissolved in DMSO at 10 mM concentration according to manufacturer’s instruction. Simultaneously, other wells were treated with 0.05% DMSO to be used as vehicle. All primary antibodies used in this study are presented in table 1. Secondary antibodies used for immunofluorescence studies were coupled to Alexa-Fluor 488 or Alexa-Fluor 546 (Molecular Probes); or to Cy3 or Cy5 (Jackson ImmunoResearch Laboratories). Secondary antibodies used for Western blotting were either horseradish peroxidase (HRP)-coupled anti-rabbit-IgG polyclonal antibodies (Jackson ImmunoResearch Laboratories) or HRP goat anti-mouse-IgG antibodies (Millipore). To visualize NMJ for immunofluorescence studies, we used α-Bungarotoxin at 5 µg/mL conjugates with either Alexa-Fluor 488 (Molecular Probes) and DAPI (D9542; Sigma-Aldrich) was used to stain nuclear DNA. To visualize and quantify proteins on Western blot, we used 2,2,2-Trichloroethanol (TCE, Sigma, #T54801) (Anand Chopra 2019 Scientific reports). For C2C12 transfection and muscle fiber electroporation experiments, paxillin (#15233), HDAC6-wt (#36188), HDAC6-ΔDC (#36189), Tub-wt (#64060) and TubK40Q (#32912) plasmids were obtained from Addgene. For knockdown of paxillin in C2C12 cells, shPXN plasmid came from Open Biosystems, GE Dharmacon RMM3981-201801357 (Clone ID: TRCN0000097194).

**Table 1:** Primary antibodies.

| antibody name | type | dilution |  | provider |
| --- | --- | --- | --- | --- |
|  |  | Immunostaining | Western blotting |  |
| Myosin Heavy Chain sarcomere | mouse monoclonal | 1:200 |  | DSHB, #MF20 |
| acetylated tubulin | mouse monoclonal | 1:200 | 1:2,000 | Sigma-Aldrich, clone 6-11 B-1 #T7451 |
| α-tubulin | mouse monoclonal | | 1:1,000 | Sigma-Aldrich, clone B-5-1-2, #T6074 |
| αTubulin (11H10) | rabbit monoclonal | 1:100 | | Cell signaling, #2125 |
| SMN | mouse monoclonal |  | 1:2,000 | BD Transduction Laboratories™, #610646 |
| Atrogin-1 (MAFbx) | rabbit polyclonal |  | 1:1,000 | ECM Biosciences, #AP2041 |
| β-actin | mouse monoclonal | | 1:2,000 | Sigma-Aldrich, #A5441 |
| GAPDH (HRP) | goat polyclonal |  | 1:20,000 | Abcam, #ab85760 |
| Paxillin (Y113) | rabbit monoclonal | 1:500 | 1:10,000 | Abcam, #ab32084 |
| Paxillin (5H11) | Mouse monoclonal | 1:100 |  | Invitrogen, #MA5-13356 |
| HDAC6 (D21B10) | rabbit monoclonal |  | 1:4,000 | Cell Signaling Technology, #7612 |

### Preparation of muscle, C2C12 cells homogenates and protein extraction

Muscles were collected from adult mouse hindlimbs and dissected muscles were crushed on dry ice. Muscle powder resuspended in urea/thiourea buffer [7 M urea, 2 M thiourea, 65 mM chaps, 100 mM DTT, 10 U DNase I, protease inhibitors (Complete; Roche/Sigma-Aldrich)] and protein concentration was determined using CB-X Protein Assay kit (G-Bioscience, St. Louis, MO).

After trypsination, C2C12 cells were solubilized in RIPA buffer [50 mM Tris– HCl, pH 8.0, 150 mM NaCl, 1% NP-40, 0.5% sodium deoxycholate, 0.1% SDS and protease inhibitors (Complete; Roche/Sigma-Aldrich)]. Protein concentration was determined using the BCA protein assay kit (Pierce/ThermoFisher Scientific) as per the manufacturer’s recommendations.

### Western blot

Five to twenty µg of total proteins were separated by SDS-PAGE supplemented with 0.5% TCE and transferred onto nitrocellulose or PVDF membranes. Non-specific binding was blocked with 4% bovine serum albumin (BSA, Euromedex) diluted in 1X PBS supplemented with Tween 0.1%, and membranes were incubated with primary antibodies. After thorough washing with 0.1% Tween 1X PBS, membranes were incubated with horseradish peroxidase (HRP)-conjugated secondary antibodies (Jackson Immunoresearch Laboratories/Cederlane). After additional washes, signals were revealed using ECL substrate reagents (Bio-Rad) and acquisitions were done using a ChemiDoc^TM^ MP Imaging Systems (Bio-Rad) or autoradiographed with X-Ray films (Fisher Scientific). Quantifications based on TCE membrane were performed with the Image Lab software (Bio-Rad) or FIJI software (ImageJ 2.0.0-rc-69/1.52n, National Institutes of Health, Bethesda, MD).

### mRNA quantification by RT-qPCR

Total RNA from cell cultures was extracted using Tri Reagent (TR118, Molecular Research Center). RNA separation was performed using cold chloroform followed by centrifugation for 15 mins at 4°C, 17000 rcf. Total RNA was then precipitated with 100% cold ethanol and 20 µg of glycogen (08-0111, Euromedex). Following precipitation, 3 RNA washes were performed using 70% cold ethanol and RNA concentration was then assessed by ultraviolet spectrometry nanodrop (IMPLEN N60, T62407). According to manufacturer’s recommendation, the extracted RNA from cell cultures was submitted to DNA enzymatic digestion (AM1907, Turbo DNase-free kit, Invitrogen) for 30 minutes at 37°C, followed by DNase inactivation and then cDNA synthesis from 1 µg of RNA with oligo(dT) using SuperScript III (18080085, Invitrogen). Quantitative real-time PCR was performed using Bio-Rad CFX384 with 1x SYBR Green ROX mix (AB1162B, Thermo Scientific) as a fluorescent detection and plated in triplicates with final volume of 7μl containing diluted cDNA and 100 nM primers (listed below). Relative mRNA expressions were normalized to PPIA and RPL13A. The normalized expression levels were calculated according to the ΔΔCt method to establish the relative expression ratio between experimental groups.

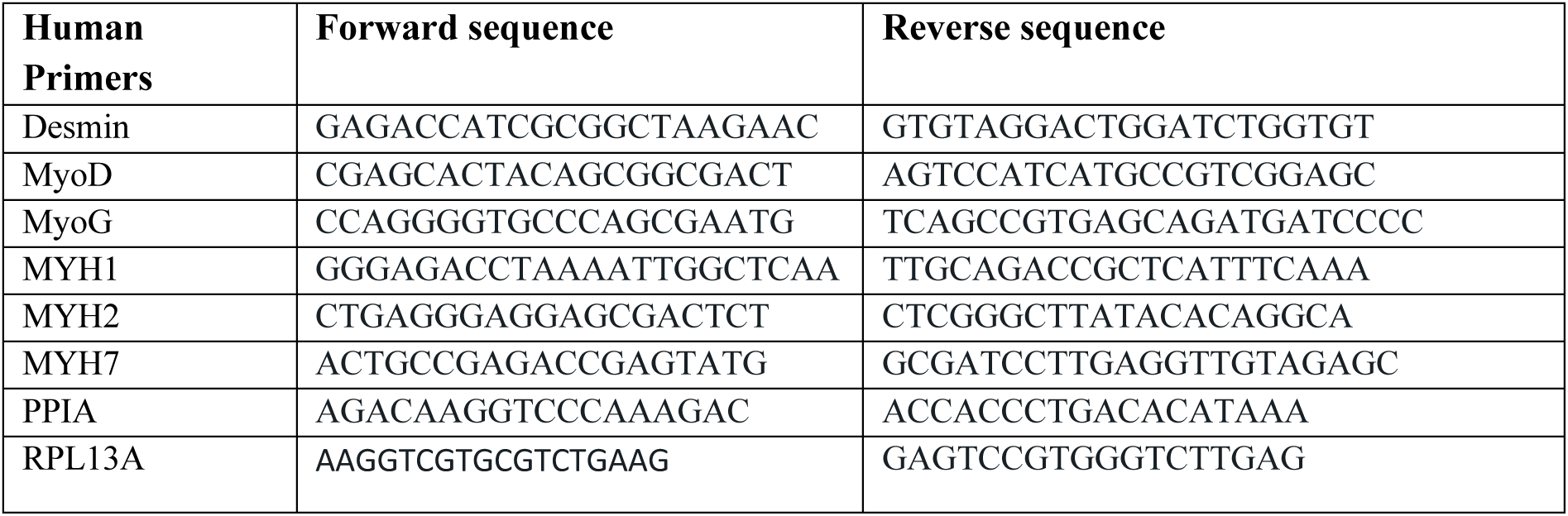

### Muscle histology and histochemistry

Frozen TA muscle samples were placed at −20 °C into the cryostat (HM 525 NX, Thermo Fisher Scientific) for at least 20 min before further processing. 10 µm thick TA muscle sections were transversally cut. TA muscle cross-sections were stained with Hematoxylin and Eosin dyes. Sections were dehydrated using 70%, 90%, and 100% ethanol solutions and washed with toluene. The sections were mounted using Permount (Fisher Scientific). Images were collected with a CMOS camera (ORCA Flash 2.8, Hamamatsu Photonics) mounted on an epifluorescence microscope system (AxioObserver Z1, ZEISS) using ZEN software (Carl Zeiss SAS) with 10 X or 20 X objective (Zeiss EC-Plan-Apo NA 0.8). For the cross-section area analysis, one section per muscle for each mouse was analyzed. Twenty percents of total fibers was measured from 5 areas dispatched randomly across the section using NIH ImageJ software.

### Immunofluorescence microscopy and image acquisition

Incubations with primary antibodies in PBS-0.1% Tween 20 were performed either at room temperature for 60 min (C2C12 or primary cells) or at 4°C overnight (isolated dissociated muscle fibers) and washed. After incubation for 1-3 hours at room temperature with fluorescent secondary antibodies, DNA nuclear were stained with DAPI for 10 min. Coverslips were mounted on microscope slides with FluorSave^TM^ reagent (Calbiochem). Images were captured at RT on either a Zeiss LSM880 microscope (Carl Zeiss) with an AiryScan1 detector equipped with a 63× 1.4-NA or a Zeiss Axio Imager M2 Zeiss (Carl Zeiss) upright microscope equipped with Plan-Apochromat equipped with either a 63× 1.4-NA or a 10× 0.45 NA with AxioCam mRm CCD detector. All images were processed with either the ZEN blue software, Zeiss AxioVision software (Zeiss, Oberkochen, Germany), Photoshop CS5 (Abobe Systems, San Jose, CA, USA) or FIJI software (ImageJ 2.0.0-rc-69/1.52n, National Institutes of Health, Bethesda, MD). Live cell imaging was performed on a Incucyte® for 48h. Images were analysed in a blinded manner by randomly renaming file names with numbers using the ‘name_randomizer’ macro in ImageJ^35^.

### Differentiation index and width myotubes – MF20 immunostaining

To evaluate the differentiation index of myotubes, cells were plated on a 6-well plate with coverslip and treatment with 5 μM TubA or 0.05% DMSO (vehicle) was launched after 2 or 3 days of differentiation for 24h. Then, cells were quickly rinsed with PBS and fixed in PFA for 10 minutes at RT and then washed 3 times for 10 minutes in PBS. Cells were incubated for 10 min at RT in PBS with 0.1% Triton X-100 for permeabilization, followed by 3 rinses in PBS for 10 mins each. Cells were then immunostained for myosin heavy chain using the MF20 mouse antibody (DSHB) diluted at 1:40 in PBS supplemented with 0.1% Tween-20 and 8% bovine serum albumin (BSA, A7906-50G, Sigma) overnight at 4°C. Following incubation, 3 washes with PBS for 10 min each were performed. Cells were then incubated with secondary antibodies. After incubation with the secondary antibody, PBS washes were performed 3 times for 10 min. Cells on the coverslips were then mounted on microscopic slides with FluorSave^TM^ reagent (Calbiochem) or Fluoromount G (Invitrogen). Images were analyzed using NIH ImageJ Software. Differentiation index was measured as the percentage of the nuclei number in MF20-positive cells relative to the total nuclei number in the field. Width myotubes result of the mean of 3 width measures in each myotube in the field.

### Statistics analyses

All statistical analyses were performed using Prism 6.0 (GraphPad Software, La Jolla, USA). Data are given as mean ± SEM. Student’s t-test was used if datasets belong to a normally distributed population with an n > 30. Otherwise, the nonparametric, two-sided U test (Mann-Whitney) was applied. Data distribution was assumed to be normal, but this was not formally tested. For a multiple factorial analysis of variance, two-way ANOVA was applied. P-values under 0.05 were considered statistically significant (shown as a single asterisk in figures, p-values below 5%); p-values under 0.01 were considered highly statistically significant (shown as two asterisks in figures, p-values below 1%); p-values under 0.001 were considered very highly statistically significant (shown as three asterisks in figures, p-values below 0.1%).

## Results

### HDAC6 regulates myotube formation

To study the role of HDAC6 in myogenesis, selective inhibitors of HDAC6 were used. Mouse C2C12 myoblasts were treated for 24 hours with either DMSO as a control or with the selective HDAC6 inhibitors, Tubastatin A (TubA) or tubacin (TBC). Total tubulin and acetylated tubulin levels were assessed either by Western blot or by immunofluorescence to confirm HDAC6 inhibition. Compared with controls, TubA or TCB similarly increased tubulin acetylation (Supp Fig. 1A-E). Proliferative myoblasts were then pre-incubated 24h with HDAC6 inhibitors (TubA or HPOB/N-hydroxy-4-(2-[(2-hydroxyethyl)(phenyl)amino]-2-oxoethyl) or a pan HDAC inhibitor (TSA/trichostatin A) and induced to differentiate for 5 days in differentiation medium. Myosin heavy chain (MHC) immunostaining and DAPI respectively allowed to measure myotube width and differentiation index. HDAC6 inhibitors and TSA all increased myotube width and differentiation index (Fig. 1A-D).

**Figure 1:**
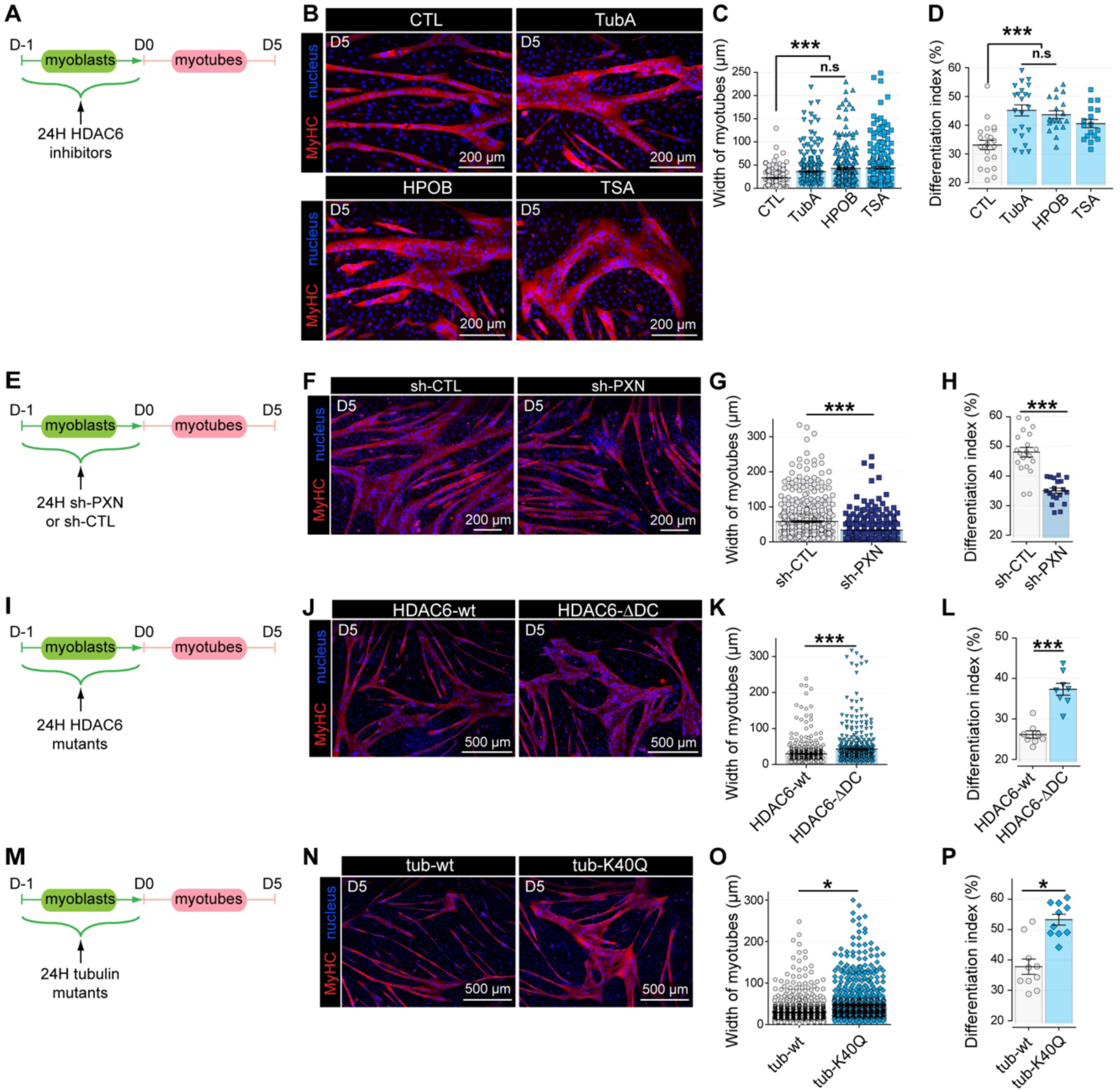
Specific inhibition of HDAC6 in myoblasts state regulates myotubes formation. (**A, E, I, M**) Schematic representations of experimental time courses. (**B**) C2C12 myoblasts were treated with the specific HDAC6 inhibitor TubA (5 µM), HBOP (5 µM), TSA (10 µM) or with DMSO (CTL; 1 µl) for 24h. (**F, J, N**) C2C12 myoblasts were transfected with either one HDAC6 mutant (HDAC6-wt; HDAC6-ΔDC) or with WT tubulin (Tub-wt) or a mutant (TubK40Q) or either with shRNA-Control (sh-CTL) or shRNA against paxillin (shPXN) for 24h. After 5 days of differentiation, myotubes were stained with an antibody against myosin heavy chain (MyHC, in red). Nuclei were labeled with DAPI (in blue). (**C, G, K, O**) Quantifications of the width myotubes were measured (in µm). (**D, H, L, P**) Quantifications of the differentiation index were measured as the percentage of the nuclei number in MF20-positive cells relative to the total nuclei number in the field. Three independent experiments for each condition. Means ± SEM. *, P < 0.05; ***, P < 0.001; Mann-Whitney U test. Scale bars: white bars in each image.

Paxillin is the only known endogenous HDAC6 inhibitor^36^. Paxillin immunostaining in C2C12 cells shows typical localization to focal adhesions as well as cytosolic staining (Supp Data Fig. 1F). To evaluate the action of paxillin on HDAC6 in myoblasts, shRNA against paxillin (sh-PXN) or control shRNA (sh-CTL) were transfected in C2C12 myoblasts. Tubulin acetylation and Paxillin levels were evaluated by Western blot and showed that Paxillin downregulation promoted tubulin deacetylation (Supp Data Fig. 1G-J). Paxillin and acetylated tubulin immunostaining confirmed this observation (Supp Data Fig. 1K-N). Paxillin action on myoblasts differentiation was then evaluated by placing myoblasts in differentiation medium 24 hours after transfection with sh-PXN or sh-CTL. Five days later, MHC immunostaining and nuclei staining allowed to evaluate myotubes width and differentiation index and to show that paxillin inhibition reduced tubulin acetylation as well as myotube size and differentiation index (Fig. 1E-H).

To further ascribe the involvement of HDAC6 into myogenesis to its tubulin deacetylase activity, deacetylase dead HDAC6 (HDAC6-ΔDC) and α-tubulin mimicking acetylation (tub-K40Q) mutants were expressed in C2C12 myoblasts. Twenty-four hours after transfection, myoblasts were placed for 5 days in differentiation medium before MHC immunostaining and nuclei staining. Overexpression of both HDAC6-ΔDC and tub-K40Q mutants’ increased myotube size and differentiation index compared with their respective controls (overexpression of wild type HDAC6 and α-tubulin) (Fig. 1I-P).

Altogether, these results indicate that HDAC6 catalytic activity and tubulin acetylation in myoblasts controls myotube formation.

### Sirt2 does not regulate tubulin acetylation in muscle cells

Both HDAC6 and sirtuin-2 (Sirt2) have been shown to deacetylate α-tubulin lysine 40^13,14^, however, the ability of Sirt2 to deacetylate tubulin in differentiated muscle cells has not been investigated. Since selective Sirt2 inhibitors have been developed^28,29^, the Sirt2 inhibitors SirReal2 and AGK2 were used to treat C2C12 myotubes. After 4 days in differentiation medium, C2C12 myotubes were incubated 24 hours in the presence of Sirt2 or HDAC6 inhibitors and tubulin acetylation was evaluated in Western blot. Conversely to TubA, neither of the Sirt2 inhibitor increased tubulin acetylation, indicating that Sirt2 activity does not significantly control tubulin acetylation in myotubes (Fig. 2A-D). Acetylated tubulin immunostaining confirmed that a 24 hours treatment with Sirt2 inhibitors did not affect tubulin acetylation in myotubes (Fig. 2E-G).

**Figure 2:**
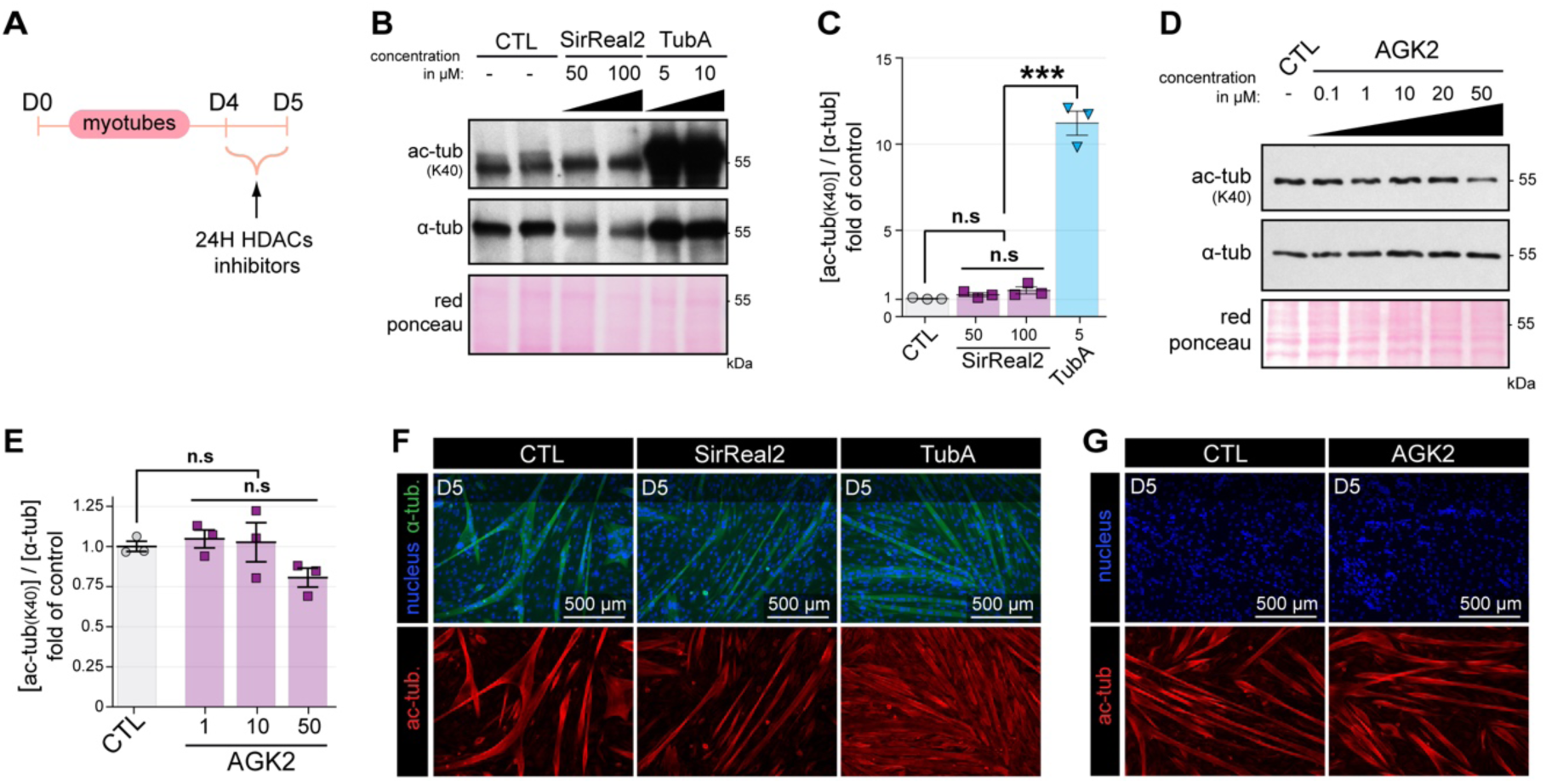
Pharmacological inhibitors of Sirt2 do not regulate tubulin deacetylation in the C2C12 cell line. **(A)** Schematic representation of the experimental time course. (**B-E**) C2C12 myotubes were treated for 24h with the specific Sirt2 inhibitors such as SirReal2 (**B, C**) and AGK2 (**D, E**). (**B, C**) Myotubes were also treated for 24h with a specific HDAC6 inhibitor, TubA (used as control). (**B, D**) Representative Western blots showing acetylated tubulin (Ac-tub) and α-tubulin (α-tub) expressions. Red ponceau was used as a loading control. (**C, E**) Quantifications of acetylated tubulin protein levels normalized with α-tubulin (n = number of independent Western blots quantified; 3). Graphs show means ± SEM. ***, P < 0.001; n.s not significant; Mann-Whitney U test. (**F, G**) Myotubes were stained with an antibody against acetylated tubulin (Ac-tub, in red), and α-tubulin (α-tub, in green). Nuclei were labeled with DAPI (in blue). Scale bars: white bars in each image.

### HDAC6 regulates myotube maturation

We then investigated the ability of HDAC6 to affect later stages of muscle differentiation. Muscle cells were treated with TubA or DMSO as a control for three days before and/or after myoblast differentiation (Fig. 3A). Different combinations of treatment were performed: DMSO before (myoblasts) and after (myotubes) differentiation (dd); DMSO on myoblasts and TubA on myotubes (dT); TubA on myoblasts and DMSO on myotubes (Td); and TubA before and after differentiation (TT). MHC immunostaining and nuclei visualization with DAPI showed that the presence of TubA, either in myoblast or in myotube or in both, increased width and differentiation index of the myotubes compared to control condition dd. No cumulative effect of TubA was observed in the TT condition compared with dT or Td conditions (Fig. 3B-D). Then, C2C12 myotubes were treated with TubA at late stage, between days 4 and 6 in differentiation medium and phase-contrast images were acquired every two hours between days 4 and 6. Alternatively, myotubes were stained with MHC antibodies and DAPI at day 6. The results indicated that inhibiting HDAC6 during the two last days of culture allow to increase myotubes width of ∼30% compared to control (Fig. 3E-H), suggesting that HDAC6 inhibition has a hypertrophic action on myotubes independent of its action on the early steps of differentiation.

**Figure 3.**
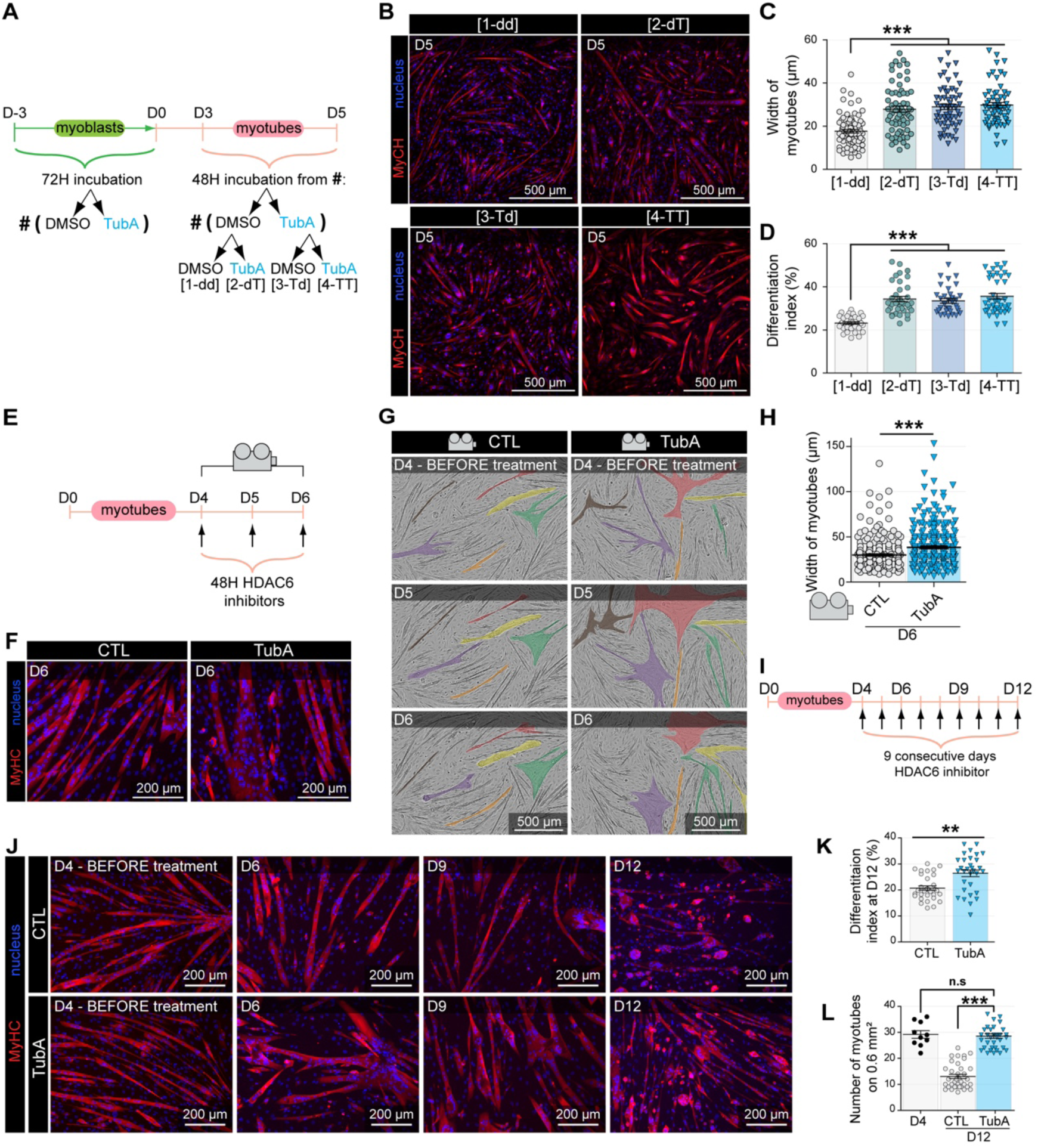
Selective inhibition of HDAC6 in C2C12 myotubes regulates their maturation. **(A)** Schematic representation of the experimental time course. (**B**) C2C12 myoblasts and myotubes were treated with the specific HDAC6 inhibitor TubA (5 µM) according the experimental time course. After 5 days of differentiation, myotubes were stained with an antibody against myosin heavy chain (MyHC, in red). Nuclei were labeled with DAPI (in blue). (**C**) Quantification of the width myotubes was measured (in µm). (**D**) Quantification of the differentiation index was measured. (**E**) Schematic representation of the experimental time lapse imaging for E–H. (**G**) Myotubes were recorded with a phase-contrast microscopy (Incucyte®) and recorded over 48 h at 30 min/frame (Video 1). Representative images are shown every 24 hours. Myotubes were colored with fictive colors. (**H**) Quantification of the width myotubes was measured (in µm). (**I**) Schematic representation of the experimental time course. (**J**) C2C12 myotubes were treated with TubA at 5 µM every day from day D4 to day D12. (**K**) Quantification of the differentiation index was measured. (**L**) Quantification of number of myotubes on 0,6 mm^2^ was measured. Means ± SEM. **, P < 0.01; ***, P < 0.001; n.s not significant; Mann-Whitney U test. Scale bars: white bars in each image.

To confirm these results, myotubes were first treated with TubA for nine consecutive days from days 4 to 12 in differentiation medium. MHC and nuclei staining revealed that in addition to the increase in differentiation index at day 12 in TubA cultures, ∼50% of myotubes had been lost in control cultures whereas in the presence of TubA the number of myotubes were preserved. These data suggest that HDAC6 inhibition favors myotube survival over time (Fig. 3I-L).

### Tubastatin A treatment improves myogenic progression in human SMA muscle cells

Despite the availability of life-saving treatments, many treated SMA patients remain disabled, mainly because of persistent muscle atrophy. To evaluate the possible benefit of selective HDAC6 inhibitors to preserve skeletal muscle in SMA patients, human primary SMA muscle cells were treated with TubA. Compared to control human primary muscle cells, the SMA muscle cells showed a reduction in SMN protein level of approximately 75% (Supp Data Fig. 2A-B). These primary cells were treated for 24 hours with TubA or DMSO and then differentiated for 3 days until myotubes were formed (Fig. 4A). MHC and DAPI staining indicated that HDAC6 inhibition increased differentiation index (P value <0.001) and myotube width (P value <0.05) (Fig. 4B-D). Similar results were obtained with control primary human muscle cells (Supp Data Fig. 2D-F). To confirm these results, immortalized human SMA muscle cells were submitted to the same protocol but this time TubA was applied during the last 24 hours of culture instead of the first day. With these cells, myotube number, width myotubes and differentiation index also increased in the presence of TubA compared to DMSO (Fig. 4E-H and Supp Data Fig. 2E-F).

**Figure 4.**
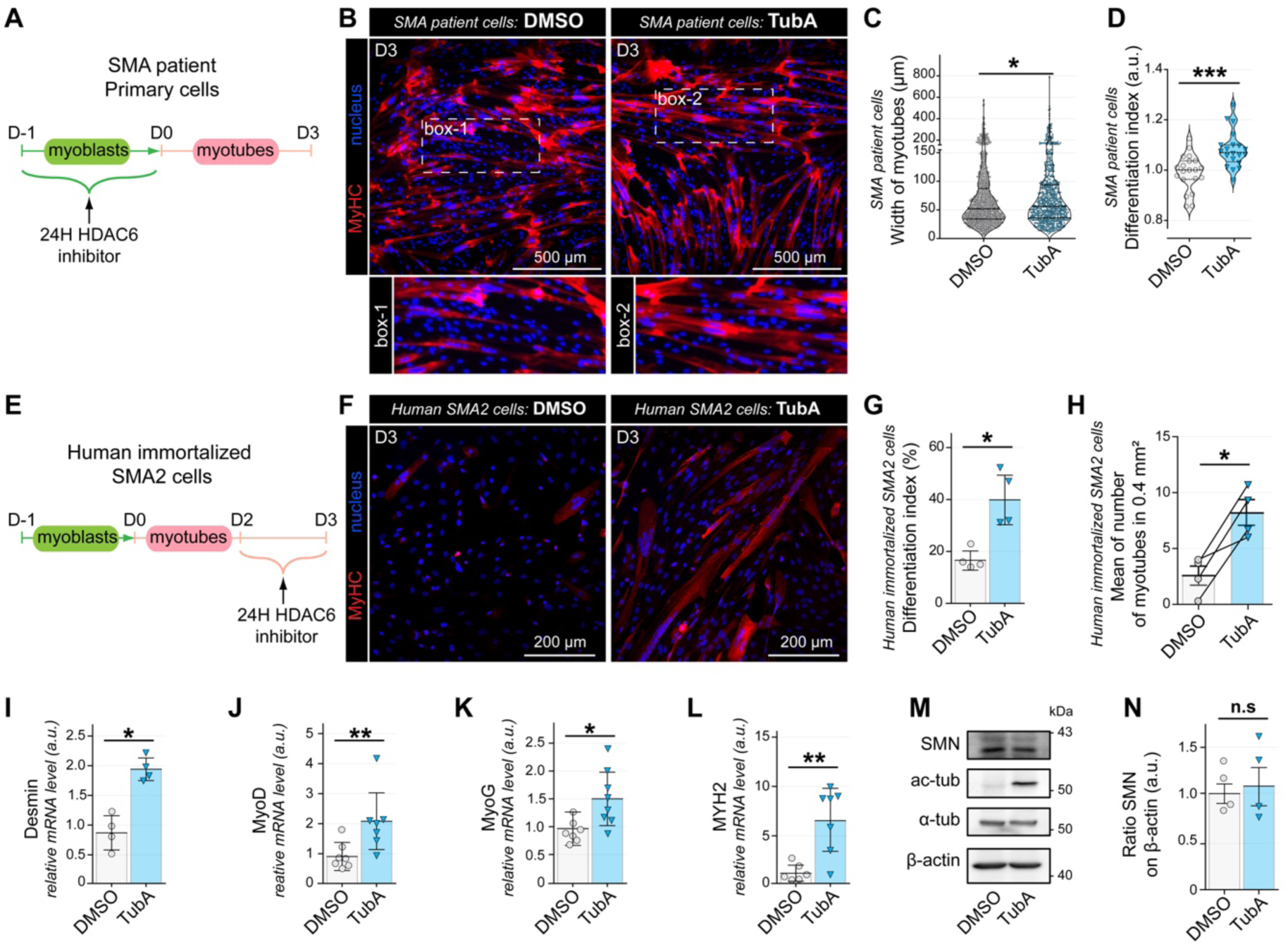
Tubastatin A treatment improves myogenic progression in human cell models of SMA. **(A)** Schematic representation of the experimental time course. (**B**) SMA patient primary myoblasts were treated with TubA (1 µM) for 24h. After 5 days of differentiation, myotubes were stained with an antibody against myosin heavy chain (MyHC, in red). Nuclei were labeled with DAPI (in blue). (**C**) Quantification of the width myotubes was measured (in µm). (**D**) Quantification of the differentiation index was measured. (**E**) Schematic representation showing the experimental time course. (**F**) Human immortalized SMA myoblasts were treated with TubA (5 µM) for 24h. After 2 days of differentiation, myotubes were stained with an antibody against myosin heavy chain (MyHC, in red). Nuclei were labeled with DAPI (in blue). (**G**) Quantification of the differentiation index was measured. (**H**) Quantification of number of myotubes on 0,4 mm^2^ was measured. mRNA levels were determined by quantitative RT-PCR in 3-day-old human SMA myotubes of Desmin (**I**), MyoD (**J**), MyoG (**K**), MYH2 (**L**). Levels of SMN, acetylated α-tubulin (ac-tub), α-tubulin (α-tub), and β-actin were performed by Western blot (**M**) and quantification of SMN protein level normalized to β-actin was evaluated (**N**). Means ± SEM. *, P < 0.05; **, P < 0.01; ***, P < 0.001; n.s not significant; Mann-Whitney U test. Scale bars: white bars in each image.

Altogether, these results demonstrate that intermittent HDAC6 inhibition can increase human SMA myotubes size and differentiation index regardless the differentiation stage at which the inhibitor is applied. We confirmed that treatment of TubA induced an 8-fold increase of tubulin acetylation in these cells (Supp Data Fig. 2G). We next evaluated the effect of TubA on the myogenic progression of the immortalized SMA muscle cell line by measuring the relative mRNA levels of myogenic markers after three days in differentiation medium and 24 hours of TubA between days 2 and 3. We quantified the mRNA levels of Desmin; a major intermediate filament, MyoD and Myogenin (MyoG); two myogenic regulatory factors (MRF) involved in myoblast determination and fusion, respectively, and Myosin Heavy Chain MYH2, which is mainly expressed in type 2a fast oxidative fibers. Compared to controls, TubA induced a 1.5-to-5-fold increase of the expression of all markers (Fig. 4I-L) whereas TubA did not change mRNA levels of MYH1 and MYH7 which are mainly expressed in fast type 2X and slow oxidative type 1 muscle fibers, respectively (Supp Fig. 2H-I). To eliminate the possibility that the beneficial effects of HDAC6 inhibition could be mediated by an increase in SMN levels, they were evaluated by western blot. As expected, TubA did not affect SMN levels in SMA muscle cells (Fig. 4M-N).

Altogether, these results indicate that HDAC6 inhibition promotes SMA muscle cells differentiation and maturation independently of SMN.

### Inhibition of HDAC6 improves muscle phenotype and protects against muscle atrophy in ASO-treated SMA-like mice

Recently, groundbreaking gene therapy treatments were recently developed, enabling children with severe forms of SMA to live. However, many SMA patients remain with a persistent and disabling muscle atrophy. Given our previous work on the role of HDAC6 in muscle atrophy^22,27^, we contemplated the possibility that HDAC6 inhibitors could be combined with SMA treatments that spare motoneurons from dying.

The results obtained above in SMA muscle cells indicated that HDAC6 inhibition was beneficial to muscle cells in the context of SMA. In order to evaluate the effect of HDAC6 inhibition *in vivo*, a population of SMA-like mice, treated by a single injection of Nusinersen-like ASOs at day 0, were treated from day 1 by a daily intraperitoneal injection of TubA (SMA-ASO-TubA) or DMSO (SMA-ASO-veh) at a dose of 10 mg/kg/day (Fig. 5A). Mice were treated either for 15 days, or all along their life. The efficiency of TubA on HDAC6 activity was assessed by measuring tubulin acetylation levels by Western blot on Tibialis anterior (TA) muscle extracts (Supp Data Fig. 3A). As expected, TubA induced a 2.8-fold increase in tubulin acetylation compared to SMA-ASO-veh muscles (Supp Data Fig. 3B). Western blot confirmed that TubA did not affect HDAC6 levels (Supp Data Fig. 3C). Together, these data indicate that TubA injection in new born mice efficiently inhibits HDAC6 deacetylase activity in muscle.

**Figure 5.**
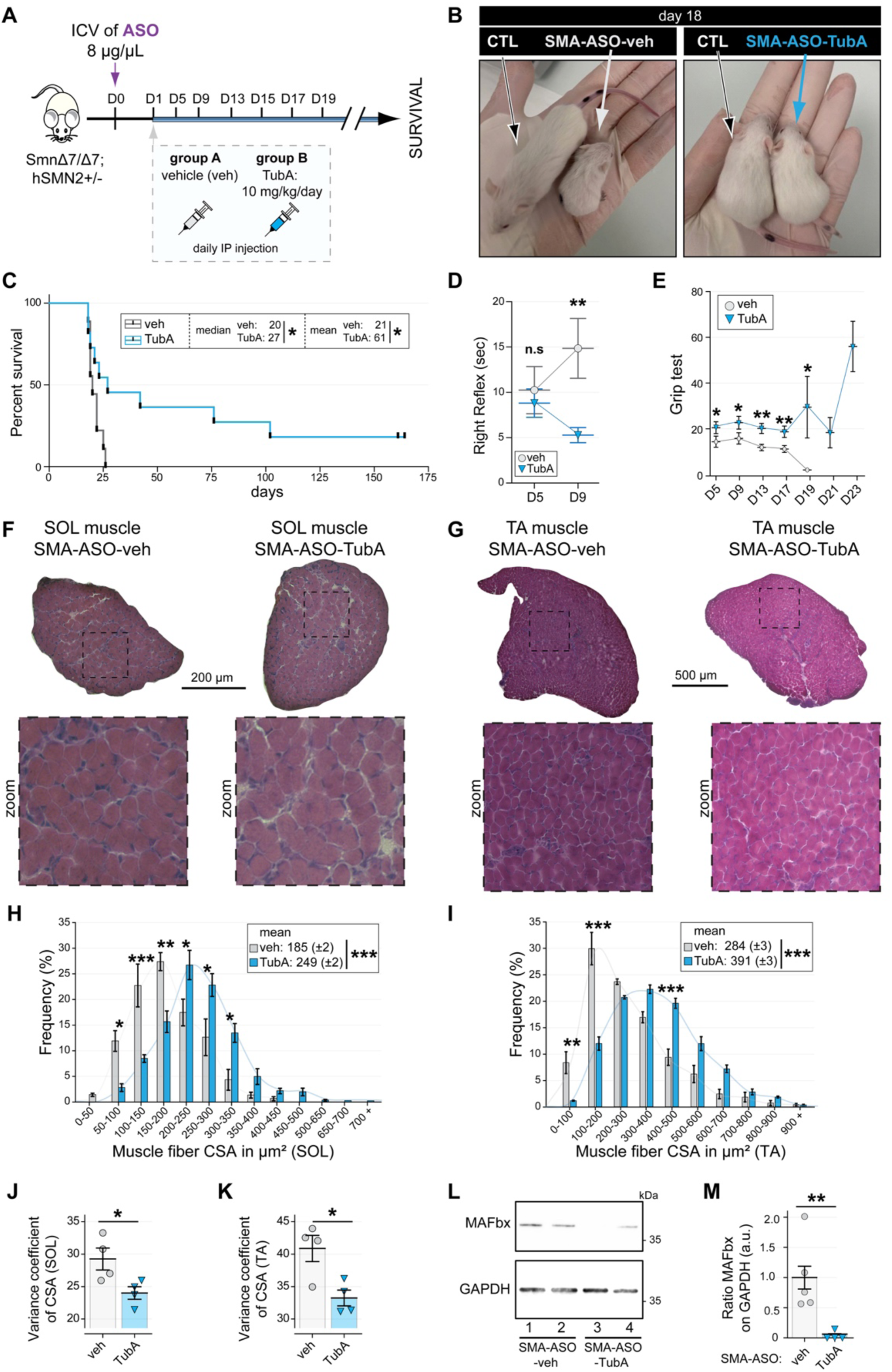
Inhibition of HDAC6 improves muscle phenotype in the SMA mouse model and protects against muscle atrophy. **(A)** Schematic representation showing the animal experimental design: All mice were injected with 8μg/μl of ASO-SMN by ICV at D0 and then divided in two groups receiving daily intraperitoneal injections of either Tubastatin A (TubA; 10mg/kg/day) or saline solution supplemented with 4% DMSO and 38.4% cyclodextrin (vehicle, veh) from post-natal day D1 to D15 for histological and molecular analysis or from D1 to the age of reaching the ethical limit, for behavioral and survival curves. (**B**) Pictures of SMA-like mice and their controls at 18 days. (**C**) Survival curves of ASO-treated severe SMA-like mice co-treated with either 10 mg/Kg/day TubA (in blue) or saline solution supplemented with 4% DMSO and 38.4% HP-β-CD as vehicles (in gray). Median survival curve equal to 20 and 27 in n=9 Veh and n=11 TubA, respectively. (**D**) The latency of the righting reflex tested at D5 (n=10 Veh, n=14 TubA) and D9 (n=12 Veh, n=14 TubA). (**E**) The grip test was measured at D5 (n=10 Veh, n=14 TubA), D9 (n=12 Veh, n=14 TubA), D13 (n=12 Veh, n=14 TubA), D17 (n=5 Veh, n=7 TubA), D19 (n=1 Veh, n=4 TubA), D21 (n=3 TubA), and D23 (n=2 TubA). Cross-section areas (CSA) of entire Soleus (SOL; **F**) and Tibialis Anterior (TA; **G**) muscles from D15 ASO-SMA-like mice treated with either TubA or veh were stained with hematoxylin and eosin. Mean CSAs of SOL (**H**) and TA (**I**) muscles are displayed above the frequency histograms in TubA (in blue) and veh (in gray). Measurements of variance coefficient in SOL (**J**) and TA (**K)** muscle fibers in TubA (in blue) and veh (in gray). n=4 mice per group, one section per muscle for each mouse was analyzed. 20% of total fibers was measured from 5 areas dispatched randomly across the section. Evaluation of levels of MAFbx protein in TA muscles, were performed by Western blots (**L**) and quantifications of MAFbx normalized to GAPDH was analyzed (**M**) (n=5 Veh, n=4 TubA). Means ± SEM. *, P < 0.05; **, P < 0.01; ***, P < 0.001; n.s not significant; Mann-Whitney U test and ANOVA test were performed. Scale bars: black bars in each image.

SMA-ASO-TubA mice were then compared to SMA-ASO-veh in terms of survival, growth and righting reflex. By day 18, SMA-ASO-TubA mice were heavier (Fig. 5B) and continued to steadily gain weight until day 75 (Supp Data Fig. 3E). TubA also significantly improved longevity with a median increase of seven days compared to SMA-ASO-veh mice, some individuals reaching more 160 days of survival (Fig. 5C) Another measure of phenotypic correction is the time of righting. Animals were placed on their back, and the time to turn and stand on their legs was recorded. After 9 days of treatment with TubA, the animals improved significantly compared to SMA-ASO-veh mice, as measured by the percentage of animals able to successfully turn around, and by the three-fold reduction of the time required to right themselves (Fig. 5D). Finally, muscle strength was evaluated daily from day 5 to day 23 using grip test in which the mice must support their weight for as long as possible. Between day 5 and day 19, TubA increased muscle strength by 1.8-fold compared to SMA-ASO-veh mice (Fig. 5E). Altogether, these data provide evidences that combining HDAC6 inhibitors to the ASO-induced significantly improve the motor behavior of the severe type SMA-like mice and significantly increase lifespan.

Then, we analyzed the muscle histology performed at day 15 in the different mouse populations and confirmed the benefit of TubA for muscle fibers. Fiber size distribution was assessed by measuring the cross-sectional area (CSA) of individual fibers in the slow oxidative soleus muscle (SOL, Fig. 5F) and the Tibialis anterior muscle (TA, Fig. 5G), of TubA- or vehicle-treated SMA-ASO mice. The muscles of SMA mice treated with the ASO were described previously as heterogenous in fiber size and containing numerous small fibers^37^. In both the TA and SOL muscles, TubA shifted fiber size distribution towards larger sizes, with a mean area of 249 vs 185 μm^2^ in SMA-ASO-veh SOL muscle and a mean area of 391 vs 284 μm^2^ in in SMA-ASO-veh TA muscle (Fig. 5H, 5I). This increase in fiber size was associated with a reduction in fiber size heterogeneity as quantified by the ∼20% reduction in the mean coefficient of variance in SMA-ASO-TubA SOL and TA muscles compared to SMA-ASO-veh muscles (Fig. 5J, 5K). Altogether, these results indicated that TubA efficiently prevented muscle atrophy in ASO-treated SMA-like. We finally evaluated whether these benefits could be correlated with a reduction in the expression of MAFbx/atrogin1, an E3 ubiquitin ligase we have previously showed to be regulated by HDAC6 in terms of activity and expression^22,26,27^. As expected, evaluation of MAFbx/atrogin1 protein levels in Western blot showed a 90% reduction in SMA-ASO-TubA TA muscle compared to SMA-ASO-veh, despite a high inter individual variability (Fig. 5L and 5M).

Altogether, these results obtained in SMA muscle cells and in ASO-treated SMA-like mice indicated that HDAC6 inhibition is beneficial to muscle cells in the context of muscle atrophy in SMA.

## Discussion

In the present study, we report for the first time that HDAC6, though the control of tubulin acetylation, is a major player of muscle fiber formation and maturation in physiological as well in pathological conditions. In spinal muscular atrophy, in which muscle fibers suffer from a persistent atrophy despite SMN-targeting treatments, HDAC6 inhibition proved to significantly improve muscle structure, including a gain in muscle fiber area and an acceleration of muscle fiber post-natal maturation. Importantly, inhibiting HDAC6 in a preclinic mouse model of severe SMA pre-treated with Nusinersen-like ASOs, resulted in the significant improvement in motor behavior and lifespan. Thus, all these data provide the first lines of evidence that HDAC6 inhibition represents a promising combinational approach to fight against muscle atrophy in SMA patients treated with SMN modulating drugs, with a potential beneficial impact on motor behavior and quality of life.

Based on the evaluation of SMA patients and animal models of SMA, SMA has long been proposed to be a multiorgan disease^38–40^. Moreover, a recent study in a SMA mouse model using AAV-driven SMN expression, has shown that restoring SMN expression in the peripheral organs provided the same increase in survival than restoring SMN in the central nervous system^41^. Of note, skeletal muscle constitutes approximately half the mass of peripheral organs. Despite currently available therapies that restore SMN expression in motoneurons, most SMA patients show a persistent and disabling muscle atrophy. Many evidences concur to show that muscle atrophy in SMA patients is not exclusively due to the denervation consecutive to motoneurons loss but also involves intrinsic muscular defects. Histological and morphometric evaluation of muscles from type I SMA fetuses revealed delayed muscle maturation^42^. Numerous studies in cultured cells and animal models also supported the notion of muscle involvement independently of motoneurons in SMA^43^. A study specifically evaluated the effect of *SMN1* inactivation in muscle precursors and showed that it caused muscle fibers defects and premature death. Importantly, this study showed that conversely to motoneurons, restoring Smn expression after the onset of the disease could reverse the phenotype^44^. Altogether these studies suggest that besides motoneurons, muscles would benefit of therapeutic intervention.

Onasemnogene abeparvovec SMA gene therapy uses AAV9 to express SMN in motoneurons. AAV9 vectors were shown to also efficiently target skeletal muscle fibers^45^. However, because of the irreversibility of motoneuron loss, SMA gene therapy has to be applied as soon as possible after birth to spare as many motoneurons as possible, and in a 2kg newborn, expressing SMN in muscle fibers would only treat 3.3% of the future muscle mass of the 60 kg adult. Until now, repeated AAV injections are not possible because of the immune response. Increasing the dose of AVV injected to treat older patients is not recommended because of the associated risk of fatal thrombotic microangiopathy. Therefore, treating newborn patients with the Nusinersen SMN2 pre-mRNA splicing modifier to allow later systemic injection of AAV vectors is not an option^46,47^. Risdiplam also targets SMN2 pre-mRNA splicing. It is administered orally and has been shown to distribute in peripheral organs and to allow a two-fold increase in the Smn protein level in muscles of SMA mice^48^. Consistently, indirect comparisons between independent clinical trials suggest that in type I SMA patients, Risdiplam sightly increases survival and motor function compared to Nusinersen^49,50^. However, no direct comparisons between Nusinersen and Risdiplam efficacy have been performed yet. Studies performed in older type 2 and 3 SMA patients do not allow to discriminate between Nusinersen and Risdiplam efficacy^49^. The SUNFISH study in type 2 and non-ambulant type 3 patients showed that 12 months of Risdiplam treatment efficiently stabilized the motor function of 59% of the patients and increased it in 16% of the patients compared to the placebo arm^51^, indicating no reversion of the muscle phenotype as observed in the muscle specific *Smn*KO mice^44^.

Altogether, currently available treatments to restore SMN expression confirm previous indications that SMA patients would benefit of complementary treatments to improve skeletal muscle function. The present study was designed to evaluate if and how HDAC6 inhibitors could be beneficial to SMA muscle cells and would provide additional benefit when combined to therapies targeting SMN. The results of the treatment of *Smn*^Δ7/Δ7^; hSMN2^+/−^ mice with a combination Nusinersen-like ASO and Tubastatin were beyond our expectations, confirming previous reports^52^ suggesting that treating skeletal muscle in the context of SMA could greatly improve motor function, health and lifespan.

The experiments performed in cultured SMA muscle cells show that HDAC6 inhibition as a cell autonomous effect on muscle cells and promotes both differentiation and myotubes maturation, which were both previously shown to be defective in SMA^53,54^. SMA mice were treated perinatally, when muscle is still actively developing, which probably explains the rapid and highly benefits of HDAC6 inhibition on muscle formation in newborn animals. An important finding reported here is that HDAC6 inhibition induced an acceleration of the muscle fiber post-natal maturation in ASO-treated SMA-like mice. We reported here for the first time and thanks to our *in vivo* and *in vitro* experiments, regardless of the specific mouse model employed, it’s reasonable to conclude that HDAC6 inhibitors can be effectively applied across various models. In this disease, this adaptability stems from two key factors: HDAC6 inhibitor’s primary effect is on enhancing muscle quality, including increased size and longevity, and the Taiwanese model, being one of the most severe, suggests that if HDAC6 inhibitor is effective in this extreme case, it’s likely to be beneficial in less severe models as well. These considerations support the potential broad applicability of HDAC6 inhibitor such as TubA across different mouse models in related research.

Moreover, although the ASO efficiently restores Smn expression in motoneurons, allowing normal function of living motoneurons at birth^55^, it cannot be excluded that HDAC6 inhibition also exerts an additional beneficial effect on motoneurons directly or indirectly, for example through the stabilization of the neuromuscular junction. Indeed, HDAC6 inhibition has been shown to be beneficial to motoneurons in the context of ALS and CMT^56,57^ and previous studies have shown the benefit for motoneurons of treatments complementary to Nusinersen^58^, two diseases in which the whole motor-unit is altered. It will therefore be interesting to design a future study aiming at analyzing in detail the effect of HDAC6 inhibition, either alone or in combination with Smn-restoring treatments, on the spinal cords of SMA mice and on cultures of SMA motoneurons.

Several treatments in combination with Nusinersen-like ASO in SMA-like mice, have proved their efficacy in synergically improve muscular symptoms. These treatments mainly included compounds that target SMN expression in the spinal cord and generally result in more SMN expression in the motoneurons^59–61^. Noteworthy, Zhou et al reported the benefits of targeting myostatin activity in the skeletal muscles of 25-mer-morpholino-treated SMA-like mice by AAV-driven inhibitors^52^. Thus, the present study reports the first use of small molecules that, tested in vivo in a preclinical model of SMA treated with Nusinersen-like ASOs, efficiently improves muscle structure, independently of SMN expression, thus paving the way for a new combinational therapy that would improve patient quality of life.

## Author contributions

AO, FC, and LS originated the project and obtained grant funding. AO, RS, FC, and LS designed its conceptualization and experiments. AO, RS, LC, ACD, LW, EB, ZC, DS and PL prepared the reagents and performed the experiments. AO, RS, LC, ACD, LW, EB, ZC, YGG, DS, CV, PL, FC and LS analyzed and interpreted the data. AO, RS, FC and LC wrote the manuscript (original draft; writing). AO, RS, LC, ACD, LW, EB, ZC, YGG, DS, CV, PL, FC and LS reviewed the manuscript and provided comments and edits.

## Acknowledgements

We acknowledge the contribution of the animal facility (U1124-T3S) of the Université Paris Cité. We thank the CIQLE microscopy facilities of University of Lyon for access to microscopes.

## Consent for publication

All authors approved this manuscript for publication.

## Data availability

Materials described in the manuscript. All relevant raw data, will be freely available to any researcher. The datasets generated during and/or analyzed during the current study are available from corresponding authors on reasonable request.

---

**Supp Data 1.**
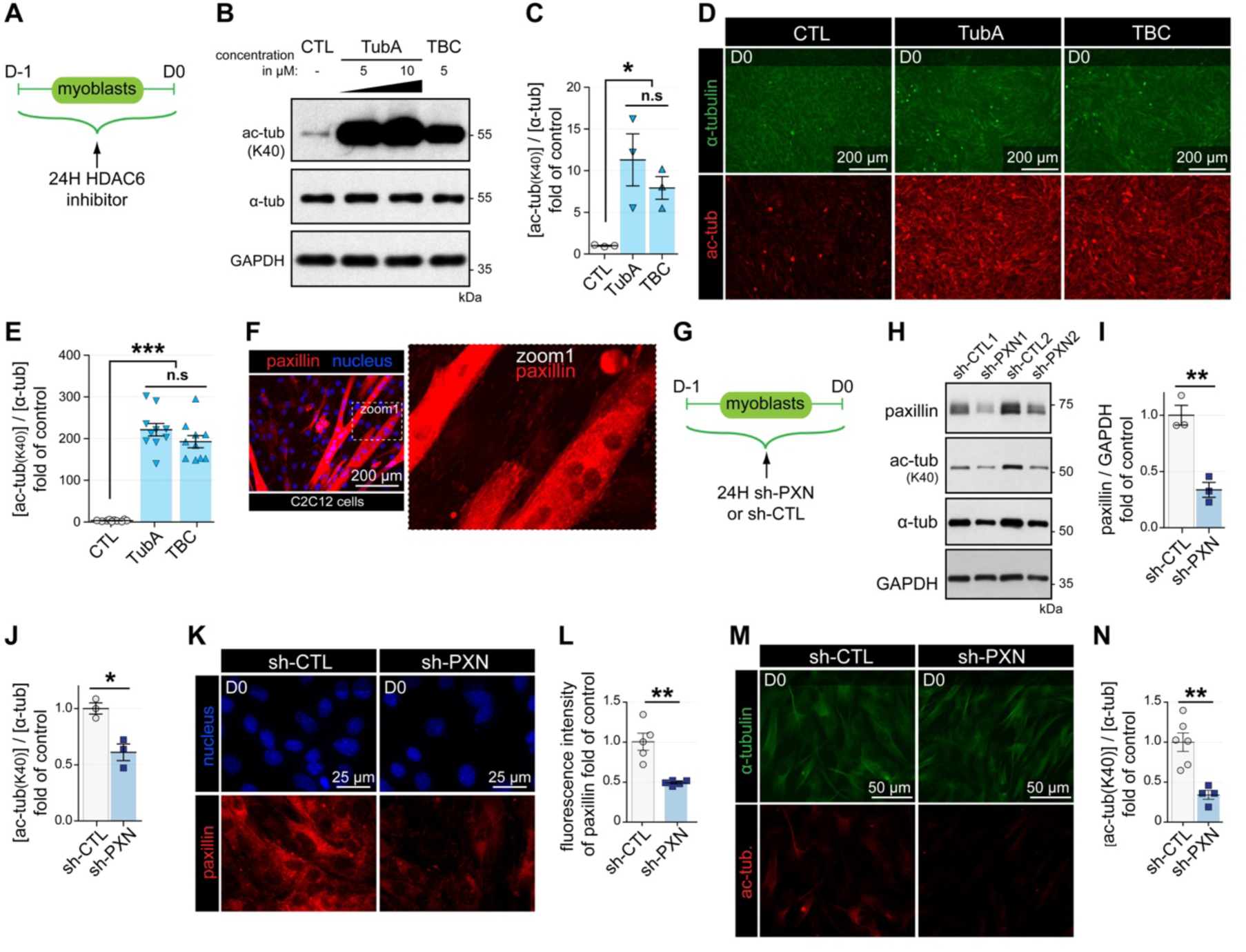
(**A**) Schematic representation of the experimental time course. (**B-F**) C2C12 myoblasts were treated for 24h with specific HDAC6 inhibitors such as TubA and tubacin (TBC). (**B**) Representative Western blots showing acetylated tubulin (Ac-tub) and α-tubulin (α-tub) expressions. GAPDH was used as a loading control. (**C**) Western blot quantifications of acetylated tubulin protein levels normalized with α-tubulin. (**D**) Myoblasts were stained with an antibody against acetylated tubulin (Ac-tub, in red), and α-tubulin (in green). (**E**) Quantifications of intensity of fluorescence of acetylated tubulin normalized with intensity of fluorescence of α-tubulin. (**F**) Myotubes were stained with an antibody against paxillin (in red) and nuclei were labeled with DAPI (in blue). (**G**) Schematic representation of the experimental time course. (H-N) C2C12 myoblasts were transfected for 24h with shRNA-Control (sh-CTL) or shRNA against paxillin (shPXN). (**H**) Representative Western blots showing acetylated tubulin (Ac-tub) and α-tubulin (α-tub) expressions. GAPDH was used as a loading control. Western blot quantifications of paxillin protein levels normalized with GAPDH (**I**) and of acetylated tubulin protein levels normalized with α-tubulin (**J**). Myoblasts were stained either with an antibody against paxillin and nuclei were labeled with DAPI (in blue) (K) or with an antibody against acetylated tubulin (Ac-tub, in red), and α-tubulin (in green) (**M**). Quantifications of intensity of fluorescence of paxillin (**L**) and of acetylated tubulin normalized with intensity of fluorescence of α-tubulin (**N**). Graphs show means ± SEM. *, P < 0.05; **, P < 0.01; ***, P < 0.001; n.s, not significant; Mann-Whitney U test. Scale bars: white bars in each image.

**Supp Data 2.**
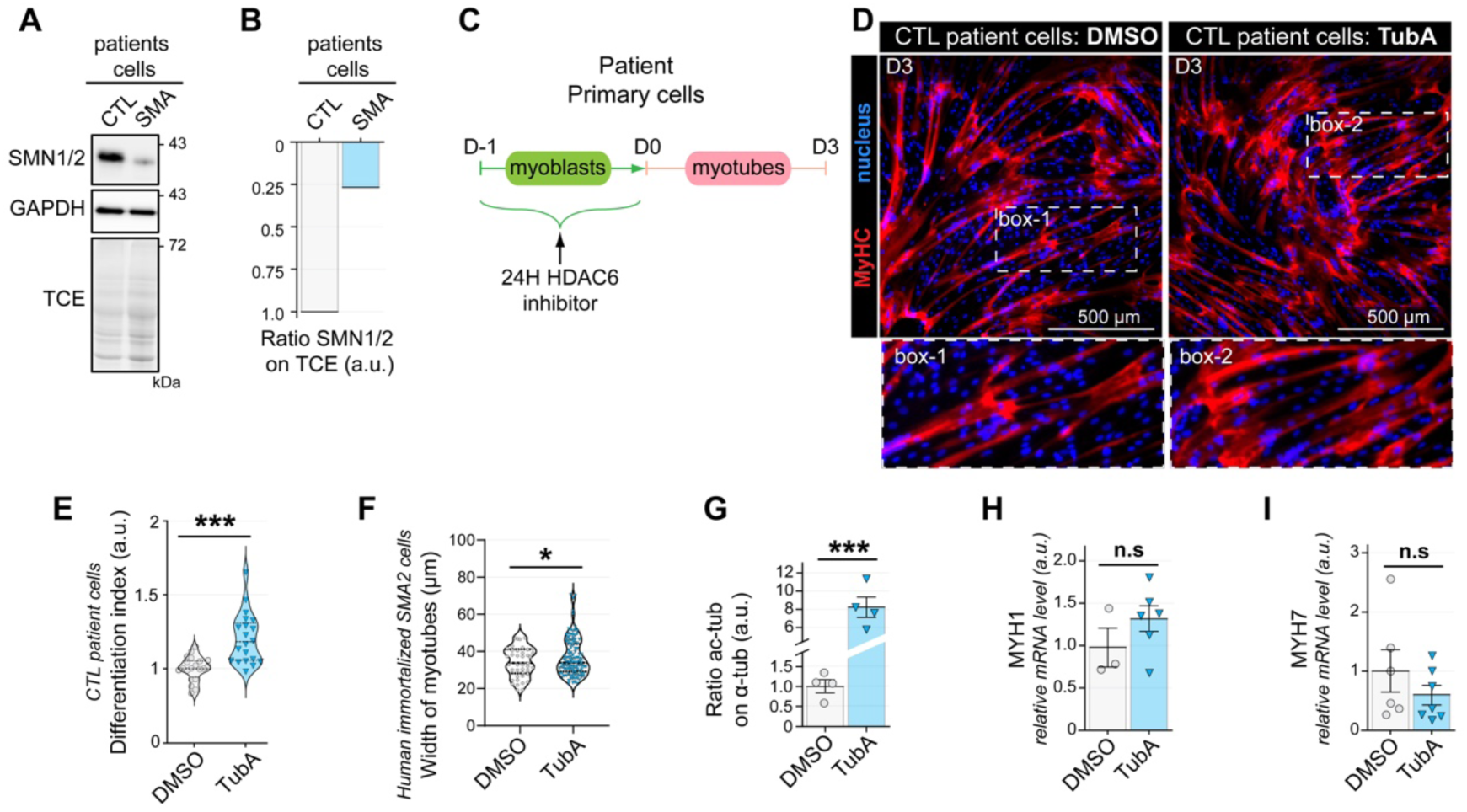
(**A**) Representative Western blots SMN1/2 expression. GAPDH and TCE were used as a loading control. (**B**) Western blot quantifications of SMN1/2 level normalized with GAPDH. (**C**) Schematic representation of the experimental time course. (**D**) CTL patient primary myoblasts were treated with TubA (1 µM) for 24h. After 5 days of differentiation, myotubes were stained with an antibody against myosin heavy chain (MyHC, in red). Nuclei were labeled with DAPI (in blue). (**E**) Quantification of the differentiation index was measured. (**F**) Quantification of the width myotubes was measured (in µm). (**G**) Quantifications on patient primary cells of acetylated tubulin protein levels normalized with α-tubulin in human immortalized SMA cells. mRNA levels were determined by quantitative RT-PCR in 3-day-old human SMA myotubes of MYH1 (**H**), MYH7 (**I**). Graphs show means ± SEM. *, P < 0.05; ***, P < 0.001; n.s, not significant; Mann-Whitney U test. Scale bars: white bars in each image.

**Supp Data 3.**
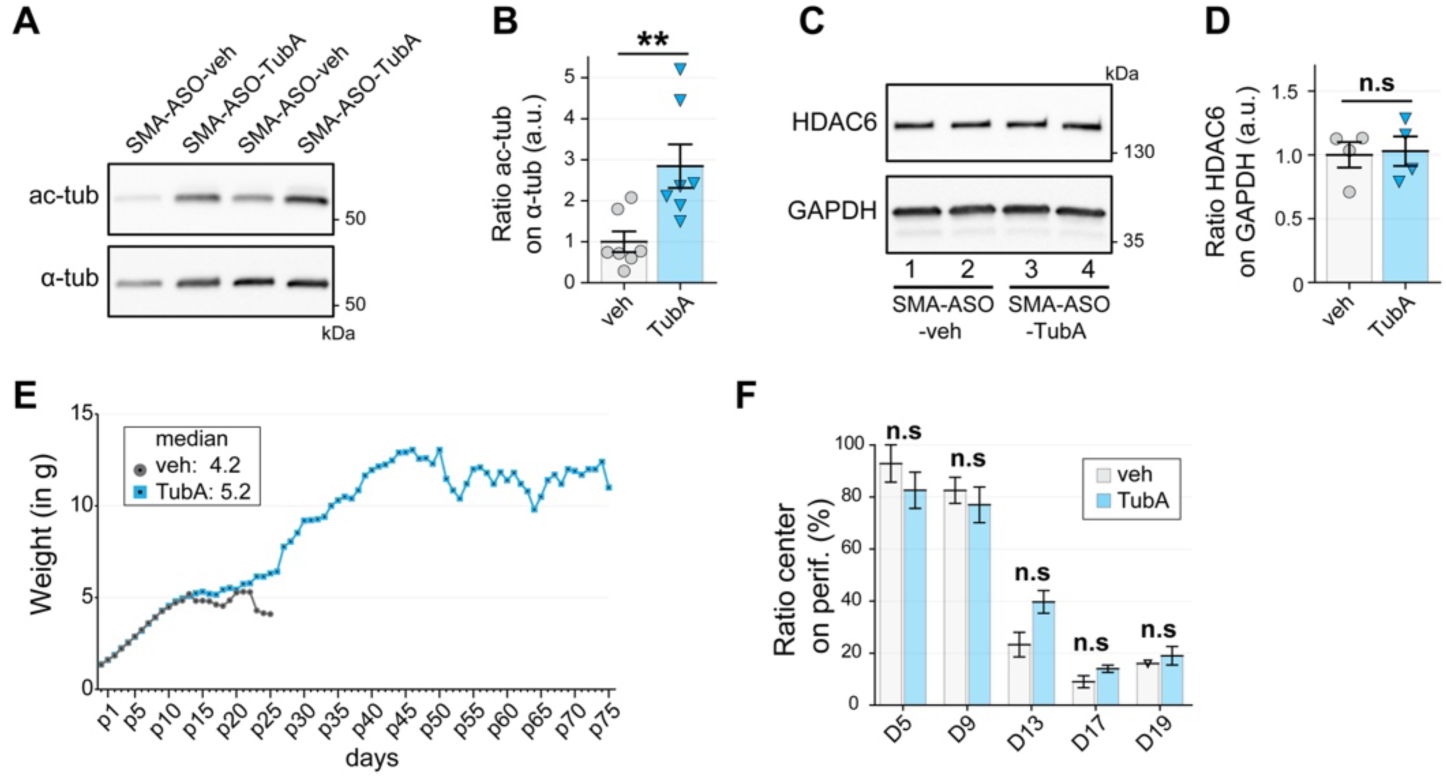
(A) Representative Western blots showing acetylated tubulin (Ac-tub) and α-tubulin (α-tub) expressions. (**B**) Western blot quantifications of acetylated tubulin protein levels normalized with α-tubulin. (**C**) Representative Western blots showing HDAC6 expression. GAPDH was used as a loading control. (**D**) Western blot quantifications of HDAC6 level normalized with GAPDH. (**E**) Growth curves of SMA-ASO-TubA and SMA-ASO-veh mice. (**F**) Open-field behavior. Time spent in the center compared to periphery areas. Graphs show means ± SEM. **, P < 0.01; n.s, not significant; Mann-Whitney U test.

## Supporting information

Video 1

